# Adaptation, spread and transmission of SARS-CoV-2 in farmed minks and related humans in the Netherlands

**DOI:** 10.1101/2021.07.13.452160

**Authors:** Lu Lu, Reina S. Sikkema, Francisca C. Velkers, David F. Nieuwenhuijse, Egil A.J. Fischer, Paola A. Meijer, Noortje Bouwmeester-Vincken, Ariene Rietveld, Marjolijn C.A. Wegdam-Blans, Paulien Tolsma, Marco Koppelman, Lidwien A.M. Smit, Renate W. Hakze-van der Honing, Wim H. M. van der Poel, Arco N. van der Spek, Marcel A. H. Spierenburg, Robert Jan Molenaar, Jan de Rond, Marieke Augustijn, Mark Woolhouse, J. Arjan Stegeman, Samantha Lycett, Bas B. Oude Munnink, Marion P. G. Koopmans

## Abstract

In the first wave of the COVID-19 pandemic (April 2020), SARS-CoV-2 was detected in farmed minks and genomic sequencing was performed on mink farms and farm personnel. Here, we describe the outbreak and use sequence data with Bayesian phylodynamic methods to explore SARS-CoV-2 transmission in minks and related humans on farms. High number of farm infections (68/126) in minks and farm related personnel (>50% of farms) were detected, with limited spread to the general human population. Three of five initial introductions of SARS-CoV-2 lead to subsequent spread between mink farms until November 2020. The largest cluster acquired a mutation in the receptor binding domain of the Spike protein (position 486), evolved faster and spread more widely and longer. Movement of people and distance between farms were statistically significant predictors of virus dispersal between farms. Our study provides novel insights into SARS-CoV-2 transmission between mink farms and highlights the importance of combing genetic information with epidemiological information at the animal-human interface.

## Introduction

Since the initial cluster of cases reported in Wuhan, China, SARS-CoV-2 is predominantly transmitted between people, with occasional examples of transmission between humans and animals. An expanding range of animals has been found to be susceptible and natural infections have been documented particularly in carnivores, including dogs, cats, lions and tigers, otters and ferrets, which were in contact with infected humans ^1,2^. Infections have not been detected in most common livestock species, but multiple countries have reported SARS-CoV-2 in farmed minks to the World Organisation for Animal Health (OIE) (https://wahis.oie.int/#/dashboards/country-or-disease-dashboard).

In the Netherlands, SARS-CoV-2 was first detected in farmed minks in late April with signs of respiratory symptoms and increased mortality^3^. An in-depth One Health investigation, combining whole genome sequencing (WGS) with epidemiological information, was conducted in response to the outbreaks in mink farms. The findings of the initial investigation between April and June highlighted that mink sequences from the first 16 farms grouped into 5 different clusters. Based on these genetic signatures, it was shown that people working on the farm were infected with mink strains rather than strains circulating among humans in the same community, providing evidence of animal to human transmission of SARS-CoV-2 within mink farms ^4^. Three of the 5 different clusters continued spreading and in total 68 out of 126 mink farms in the Netherlands were diagnosed with SARS-CoV-2 infections between April and November 2020. From January 2021 onwards all fur farming was banned in the Netherlands. To date, the mode and mechanism of most farm-to-farm transmissions have remained unknown. Phylodynamic analyses of whole genome viral sequences from mink farms and associated human cases combined with epidemiological data can help to address specific epidemiological and outbreak control questions.

In this study, we describe an in-depth molecular epidemiological analysis of the outbreak in 68 mink farms in the Netherlands, as well as related humans on these mink farms. We used Bayesian phylodynamic methods to gain more insight in the timing of SARS-CoV-2 introductions and the patterns of farm-to-farm transmission. Specifically, we explored the approximate time of onset for the different mink farm clusters and we compared the rate of evolution and population dynamics between mink clusters with the rate of evolution in the human population. Further, we have quantified the virus transmission patterns between different farms and identified farms which are more likely to be the donors of such transmissions; finally, we tried to infer the possible predictors that may drive the transmissions between farms.

## Results

### SARS-CoV-2 infections in mink farms in the Netherlands

In total 68 (farm IDs:NB1 to NB68) of 126 mink farms in the Netherlands were diagnosed with SARS-CoV-2 between the 24^th^ of April and the 4^th^ of November, and these farms were culled within 0-6 days after sampling from NB8 onwards (mean 2, median 1) (Figure 1a and b). Control measures were implemented immediately after the first infected farms were detected and included culling of infected farms from June onwards. All mink farms were subjected to a ban on transport of animals, animal materials, visitors and implementation of strict hygiene protocols and animal surveillance programs for early detection (Figure 1a).

**Figure 1.**
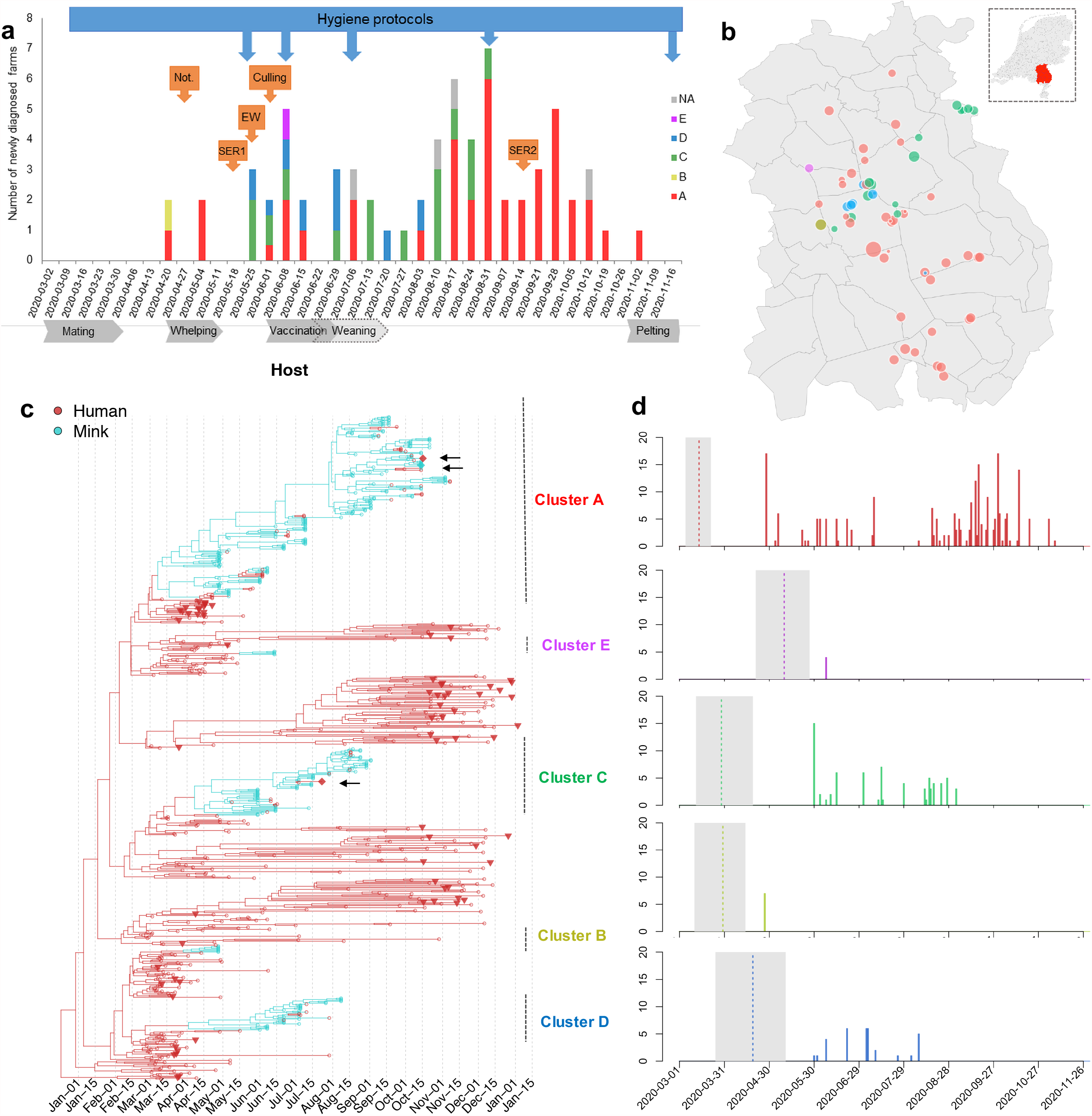
Distinct Clusters of SARS-Cov-2 circulating in mink farms in Netherlands. **a** Overview of SARS-CoV-2 outbreaks on mink farms in the Netherlands in relation to implementation of control measures and the mink farm cycle. The diagnosed farms per week are colored based on cluster. One farm in week 2020-06-01 is indicated as half A/half D as both clusters were found. The blue arrows above the graph point to the starting week of implementation of more strict hygiene protocols with regard to people working on or visiting farms. Orange arrows point to the start of other control measures including obligation for notification of clinical signs and mortality (Not.), first and second serological screening (SER1 and 2), early warning system with weekly sending in of carcasses (EW) and culling of infected farms. Below the graph important periods in the farm mink cycle are indicated. These include generally the mating season (March), whelping (April/May), vaccination (June) and weaning (June and July). Also, the start of the pelting season is shown. **b** The location of sequences isolated from each mink farm. The locations of farms on the map have been jittered for privacy reasons. **c** Time-scaled MCC tree of SARS-Cov2 sequences isolated from humans and minks in the Netherlands (n=673). Humans in red and minks in green, the subsampled human samples (n=72) isolated from the same 4-digit post code are highlighted as triangle, and 3 samples (1 escaped mink and 2 unrelated human sequences) which fell within mink clusters are highlighted as diamond and indicated by arrows. Clusters of sequences from minks and the lineages are indicated on the right. **d** The number of samples in time for each cluster. The estimated TMRCAs of each cluster are indicated via dotted line (mean) and grey shade (95% HPD intervals).

Most SARS-CoV-2 positive farms were located in a mink farm dense area in the south-east of the Netherlands with 43 farms positive in the province North Brabant and 23 out of 68 farms positive in Limburg (Figure 1b). Two farms were located in the province Gelderland, bordering on another mink farm dense area. Up to July, on average 1.73 farms (median 2) were diagnosed per week. Despite implemented control measures, and cessation of activities involving handling of the minks and employing additional staff after the weaning period in July, the weekly number of farms diagnosed increased in August and September to 3.75 (median 3.5), after which it declined to 1.34 (median 1) in October and November (Figure 1b).

At 41/68 mink farms, employees were confirmed SARS-CoV-2 positive by RT-PCR. Sequences belonging to all five mink clusters were identified from these human samples on each farm, varying from 55% of farms in cluster A (22/40) to 100% in cluster B and E (1/1). On 31 out of 41 farms, the sampling date of the human positives was after the date their minks reported positive while for two farms the human sampling dates were unknown. In three out of eight farms, workers tested positive over one week before their animals were reported to be SARS-CoV-2 positive.

Between 24^th^ April 2020 to 4^th^ November 2020, we have obtained full genome sequences of 295 minks from 64 out of the 68 mink farms. No genomes were available from 4 farms (NB22, NB30, NB37 and NB66). From 57 out of 102 human positives directly linked to 27 farms, a full sequence was obtained.

### Introductions and ongoing spreading clusters in mink farms

To look at the transmission of SARS-CoV-2 in mink farms in the Netherlands, we included full length genomes of SARS-CoV-2 from humans and animals infected on mink farms, and representative SARS-CoV-2 genomes from COVID-19 cases from the general human population of the Netherlands (n=673) to perform a time resolved phylogeographic analysis using BEAST (Figure 1c). The 5 distinct mink farm sequence clusters (A-E) were derived from 4 lineages B.1.8 (Cluster A), B.11 (Cluster B and D), B.1.22 (Cluster C) and B.1.5 (Cluster E) which have been dominantly circulating in the general human population in the Netherlands according to the Pango-lineage descriptions^5^(version on 1^st^ of April 2021).

The largest cluster found on mink farms is the so-called Cluster A, which contains 195 sequences isolated from approximately 60% of the infected mink farms (n=40) across 15 municipalities in three provinces sampled between early April to mid-October 2020 (Figure 1b). Cluster C and D have been sampled from fewer farms and circulated for shorter time periods: Cluster C viruses were isolated from 15 mink farms between late May to early September while Cluster D viruses were isolated from 8 mink farms from late May till early August. In comparison, Cluster B and Cluster E have only been identified on one farm (NB2 and NB11, respectively) in the early stage of the epizootic with no subsequent spread. The majority of farms were located within 3 km of each other, but not all neighboring farms were infected with a virus from the same cluster (Figure 1b).

Seventeen human SARS-CoV-2 sequences from farm NB1-16 between April-May have been described previously ^4^, and here we report another 35 human sequences of mink farm employees in the period June-November (tips in red in Figure 1c). All human sequences were part of the mink-related Clusters A, C and D indicating ongoing transmission between minks and humans (or vice-versa) within the three clusters. All but one of the human sequences were part of the same cluster and were closely related to the sequences of the minks on the same farm. One human sequence of a Cluster C farm (NB 24) belonged to Cluster D, which could be explained by the fact that this employee assisted in the culling of minks at another farm, where minks were infected with a Cluster D virus.

Interestingly, unique clusters were found on the majority of infected farms, only in one farm two different clusters were found: NB8 (infected viruses belong to both Cluster A and D in early June). It is therefore likely this farm was exposed to two sources of viruses.

We estimated the evolution rates of SARS-CoV-2 in mink populations in the Netherlands by using relaxed clock models, with a mean clock rate of 7.9×10^−4^ subst/site/year with 95% highest posterior density (HPD) (7.2 ×10^−4^, 8.4 ×10^−4^). The approximate times for the ancestral jumps from humans to minks were between mid-March (Cluster A, B and C) to late-April (Cluster D and Cluster E) (Figure 1d). Three clusters (A, C and D) had ongoing spread to more farms from June to November after the initial investigations of the 16 farms between April to June 2020.

The last infected farm was detected on the 4^th^ of November, after which no new infections were detected (Figure 1a).

### Spill-over into local community and limited onward transmission

In total, 218 sequences isolated from randomly selected patients from 31 postal codes, in the region of SARS-CoV-2 positive mink farms were obtained in period 4^th^ March 2020 to ^4th^ January 2021, to assess possible spill-over to the local community. In addition, all sequences submitted to GISAID from the Netherlands until the 4^th^ of January were included in the analysis.

In three separate occasions, a mink related strain, linked to clusters A and C (Figure 1c), was detected. Two out of three patients infected with a mink strain (sampling dates in July and August), lived in a province where no infected minks were reported, and they did not have direct or indirect contact with the mink farming sector. One patient was found in the regional screening in November but also did not report any mink farm contacts. After November, no human infections with mink strains have been detected (Figure 1c).

Throat swabs of the two escaped mink, caught 8 and 9 days in close proximity to two culled farms (NB58 and NB59), at 450 and 650 m distance respectively, tested positive for SARS-CoV-2 RNA. Genome sequencing was successful for one mink sample, and it belonged to Cluster A (Figure 1c).

### Specific mutations in the Spike protein in multiple mink clusters

We further explored how the specific mutations in the spike protein are associated with phylogenies by mapping 4 potential important mutations in the spike protein (L452M, Y453F, F486L, N501T) on the tree composed of the complete dataset (Figure 2). These 4 mutations are in confirmed contact residues of the viral spike protein with the ACE2 receptor ^6,7^. Within the Netherlands, these ‘mink specific’ mutations were only found in minks and employees on mink farms by the time the analysis has been performed (by 1^st^ April 2021), except for 3 samples: two sequences from unrelated humans (one with F486L, the other with both F486L and L452M, third sequence excluded due to insufficient coverage) and one sequence from an escaped mink (with F486L). However, these mutations have also been seen elsewhere in other independent lineages. For example, the F486L has been detected occasionally in humans in Ireland and Columbia, and in mink samples from the US (http://cov-glue.cvr.gla.ac.uk/#/home).

**Figure 2.**
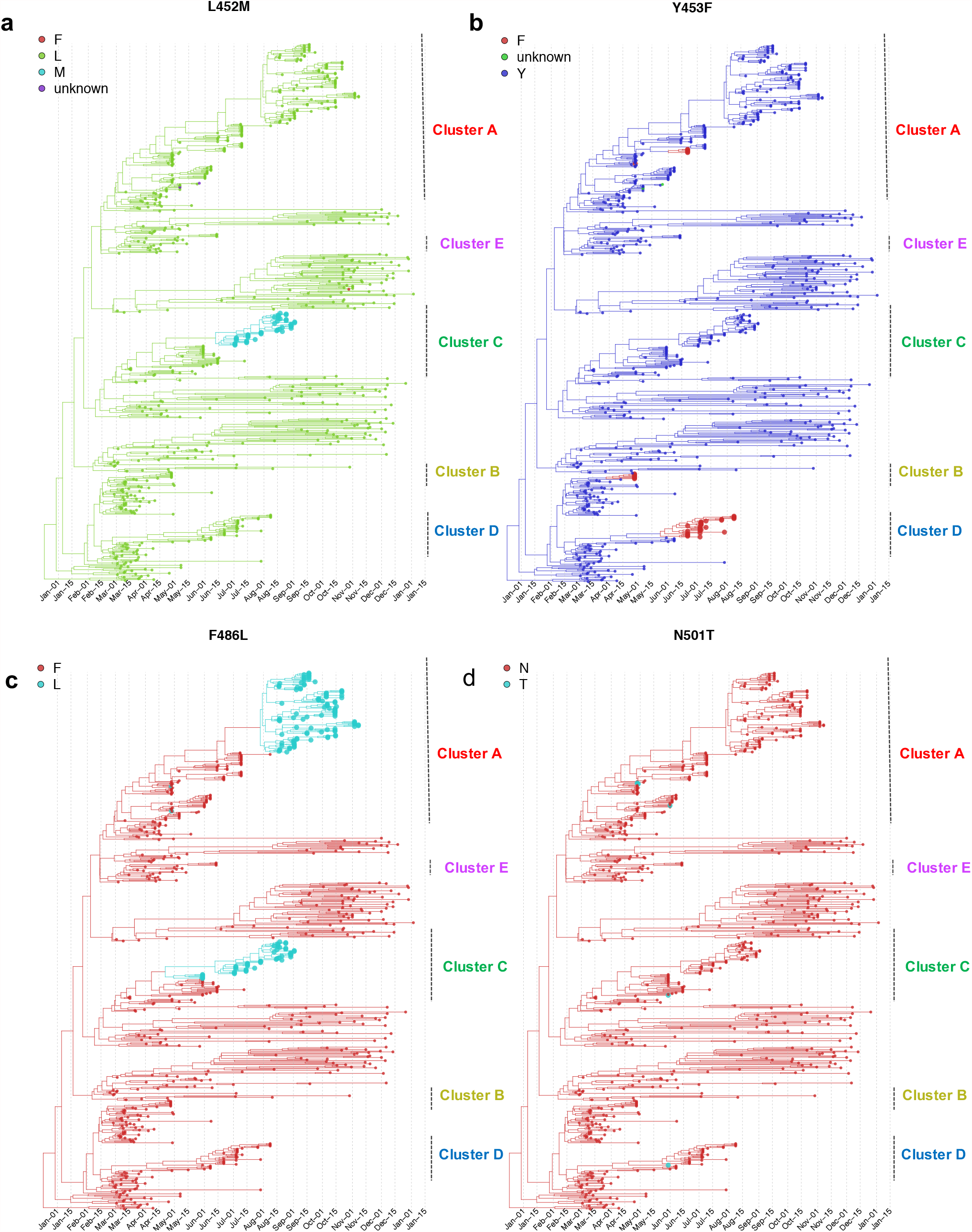
Time-scaled MCC tree of SARS-Cov2 sequences mapping with 4 mutations **a** L452M, **b** Y453F, **c** F486L and **d** N501T of the spike protein (n=673). Tips with mink specific mutations are enlarged. The phylogeny is the same as Figure 1a.

The 4 mutations have evolved in multiple clusters and in both human and mink samples from Dutch mink farms. Specifically, mutation F486L has been seen in 217 sequences from 40 mink farms that belong to 2 separate clusters (A and C), which accounted for 67% sequences and 68% sequences isolated within the cluster. Y453F has been seen in 37 sequences from 10 different farms in 3 different Clusters (A, D and E), which accounted for 3%, 82% and 100% sequences isolated within the cluster. In addition, we found the N501T mutation in only 3 mink virus sequences from 3 different farms belonging to Clusters A and D. L452M was seen in 44 sequences isolated from 9 mink farms all belonging to Cluster C (59%). N501T only appeared in a short period of the outbreak (end of April to end of May), while the others appeared in a later stage and sustained longer (F486L first appeared in two sequences in Cluster A at the end of April, then reappeared and replaced F486 in Cluster A since mid-August and in Cluster C since June, respectively); L452M appeared from early July to September and Y453F appeared from end of April to early July.

We mapped 4 types of traits (host, farm ID, province and municipality) on individual time-scaled phylogenies of Cluster A, C and D using discrete trait models. We compared the 4 individual mutations in the spike protein and the combinations of the 4 mutations on the time-scaled phylogenies of Cluster A, C and D independently. The discrete trait mapping trees of Cluster A are shown in Figure 3, with the branches and nodes colored by inferred ancestral traits. The trees for Cluster C and Cluster D are shown in Figure S1 and Figure S2. The occurrence of the mutations did not show any significant association either to host types, to farm numbers or to locations (Mann-Whitney U Test, with p>0.5).

**Figure 3.**
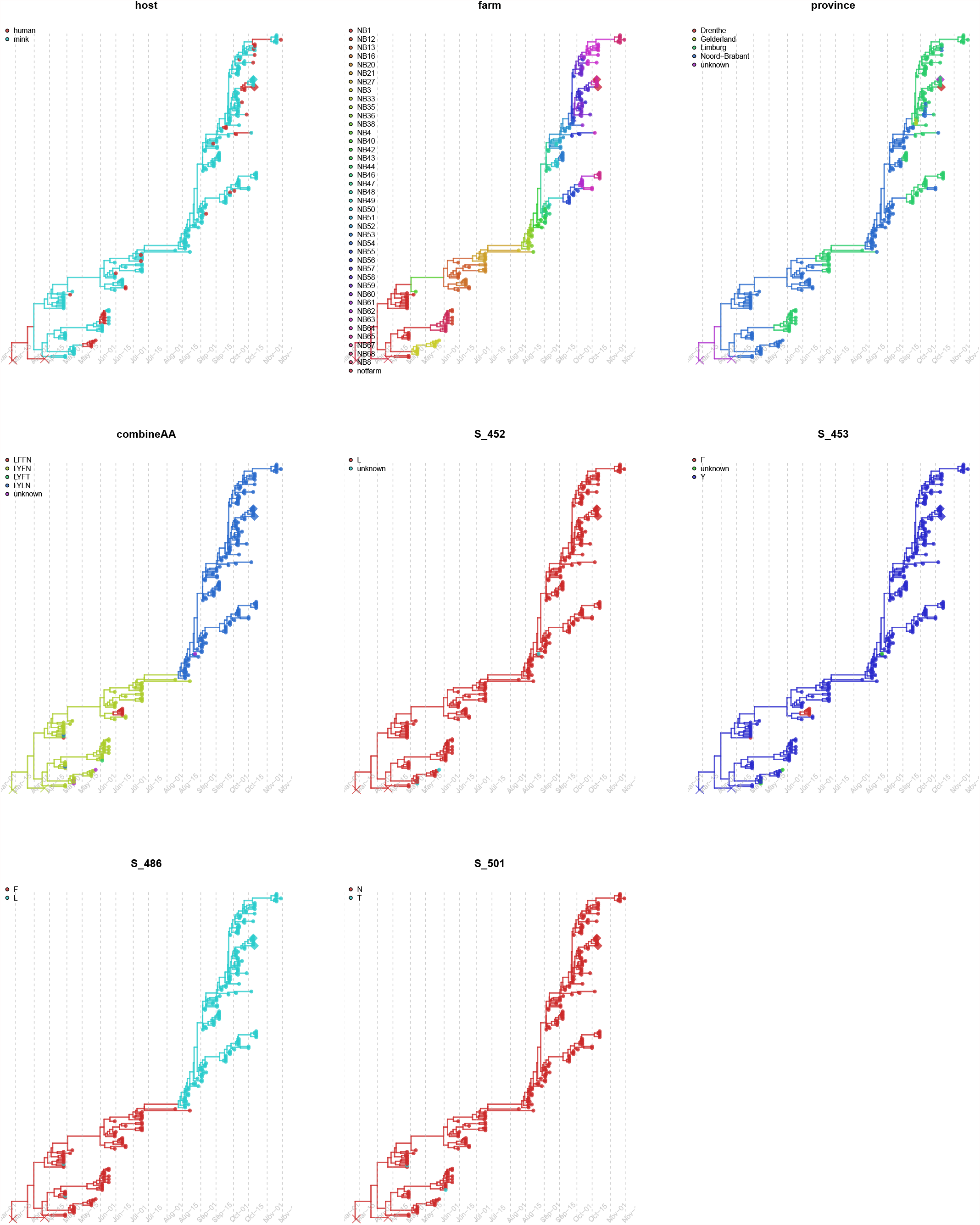
Discrete trait mapping on time-scaled phylogeny of Cluster A. Nine traits including host, farm number, province, the 4 individual mutations in the spike protein (L452M, Y453F, F486Land N501T) and the combinations of the 4 mutations are mapped on Cluster A tree using discrete trait model, with the branches and nodes are colored by inferred ancestral traits. Samples (1 escaped mink and 1unrelated human sequences) fell within mink clusters but not isolated from farms are highlighted in diamond. The outgroup human samples in the origin are cross labelled.

### Comparisons of the phylodynamics of different clusters in minks

We compared the phylodynamics of three clusters (A, C and D). The results of estimating the time to the most common recent ancestor (TMRCA), the molecular clock evolutionary rate and spatial diffusion rate (geography.clock.rate) according to available data and parameters selected are shown in Figure 4. For Cluster A, the estimated TMRCA for mink sequences is approximately in mid-March 2020 (mean 15^th^ March 2020 with 95% HPD (12^th^ March 2020, 28^th^ March 2020); Evolution rate is approximately 1.41 ×10^−3^ subst/site/year with 95% HPD (1.2 ×10^−3^, 1.75 ×10^−3^) subst/site/year. The other two clusters have slightly lower evolution rates and more recent TMRCAs, but with wider HPD intervals; overall these results are consistent with the estimations using a relax clock model on the complete data in Figure 1. Similarly, the spatial diffusion rate of Cluster A is higher (means of 2.91 ×10^−4^) than the other two clusters C and D, which have means of 1.06 ×10^−4^ and 1.34×10^−4^ (Figure 4 and Table S1). Overall, the TMRCA aligns with the epidemiological data about the emergence and detection of SARS-CoV-2 in the Netherlands. It also has a faster and wider spatial spread and higher evolutionary rate than the other clusters.

**Figure 4.**
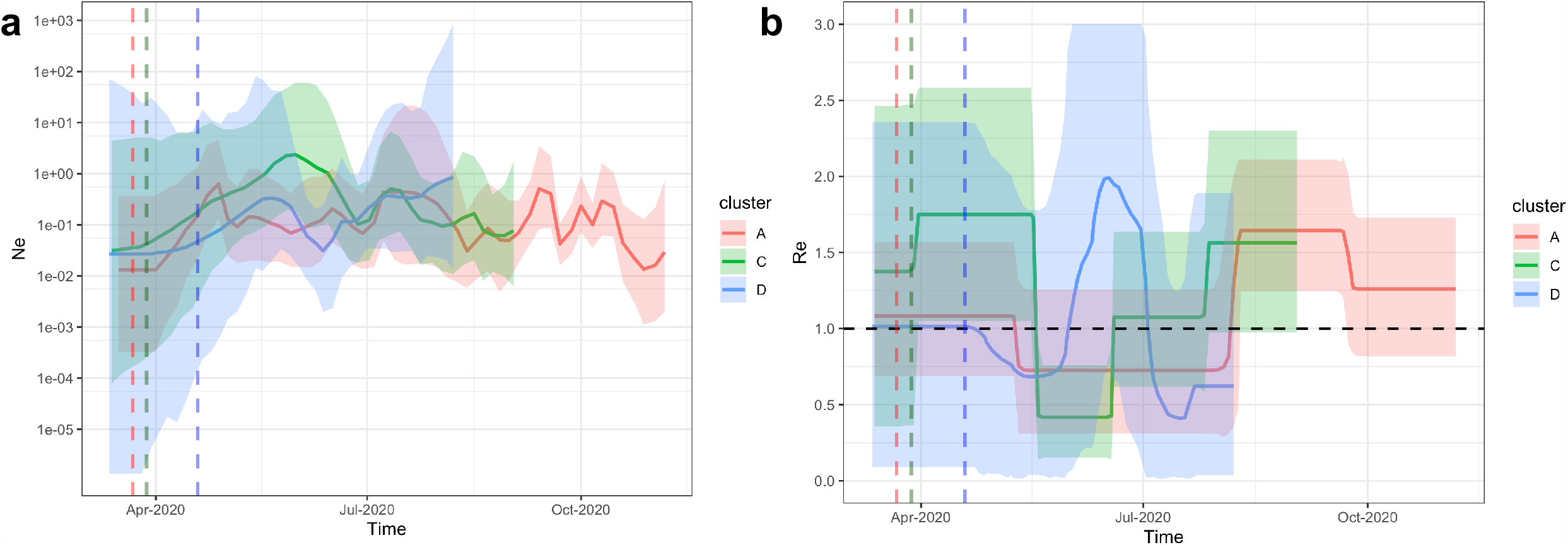
Comparisons of TMRCA, evolution rate and spatial diffusion rate of Cluster A, C and D. **a** The mean TMRCA and 95%HPD interval for each cluster. **b** The mean clock rate and 95% HPD interval for each Cluster. **c** The mean spatial diffusion rate and 95% HPD for each cluster.

We further compared the population dynamics and the transmission potential of different clusters. The estimated effective population size (Ne) and the estimated reproductive number (Re) are shown in Figure 4a and 4b. The phylodynamic Re is a relative growth rate and can be thought of as representing infection on a between-farm level rather than a between animal level given the limited number of sequences (5 on average) sampled per farm. Different patterns of Ne and Re were observed for Cluster A overtime, and for Cluster C and D. For the largest, Cluster A, the population size of mink farm sequences experienced an expansion in late March 2020 and fluctuated later on. Re for Cluster A stayed above 1 after the start of infections then decreased slightly since May 2020. The rate increased again and peaked at approximately 1.6 with 95% HPD (1.2, 2.1) since early August 2020 and dropped to 1.3 with 95% HPD (0.8, 1.7) from the end of September till November 2020. For Ne of Cluster C, a period of slight increase was observed in mid-June 2020, followed by a decline in size from June to September 2020. The Re for Cluster C stayed above 1 till May 2020 then decreased sharply (below 1) and increased again and stayed at around 1.5 with 95% HPD (1.0, 2.3) from the end of July 2020. In comparison to Cluster A and C, both Ne and Re for Cluster D have larger uncertainties (wide HPD intervals). These results are in line with the detection of SARS-CoV-2 in mink farms, few farms were infected in July 2020 while there was an increase from August 2020 onwards (Figure 1a). In addition, the timing of Re increases in the later stage coincides with the appearance of clades with mutations on Spike protein: F486L (in Cluster A and C), L452M (in Cluster C), Y453F (in Cluster D) (Figure 2, 3 and Figure 5). We observed similar results by using the multi-type birth–death model which showed a strong increase in the number of infections in clades with mutations rather than clades without mutations (Figure S3).

**Figure 5.**
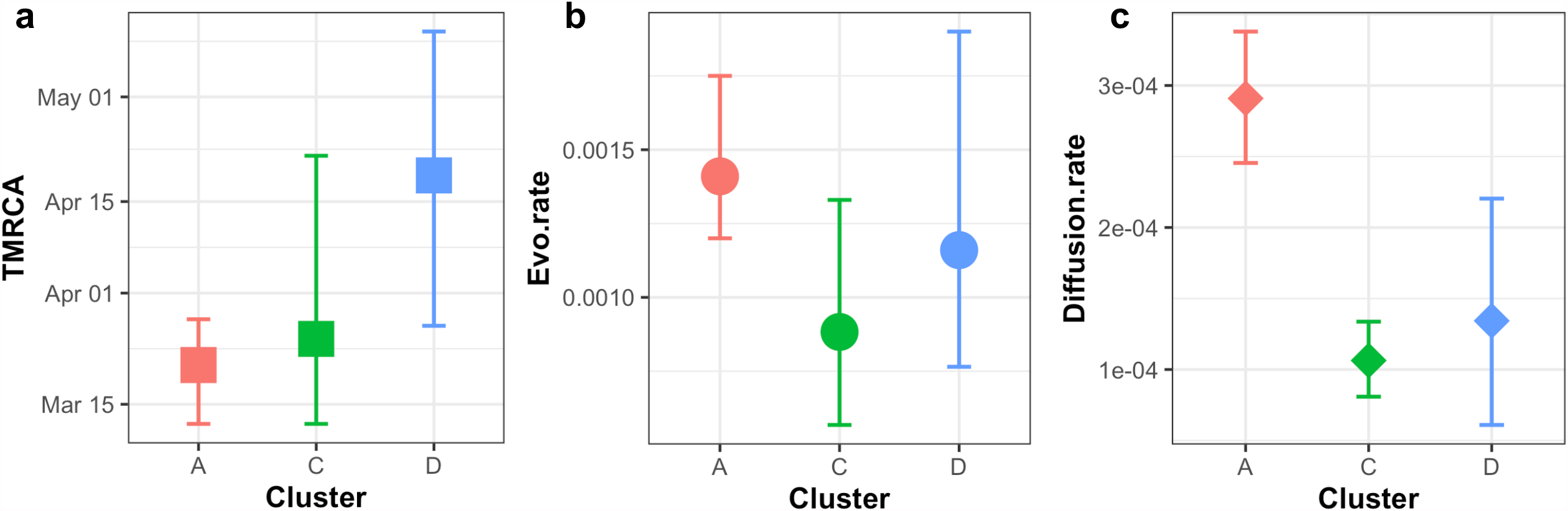
Bayesian skygrid and BDSKY analysis reveal spatiotemporal independent population dynamics of Cluster A, C and D. **a** Estimation of effective population size by skygrid analysis for Cluster A (red), C (green) and D (blue) sequences. The logarithmic effective number of infections (*Ne*) viral generation time (*t*) representing effective transmissions is plotted over time. 95% HPD intervals are plotted in lighter colors. Vertical dashed line is the mean TMRCA. **b** Estimation of Reproductive number Re by BDSKY analysis of Cluster A (red), C (green) and D (blue) sequences. The shaded portion is the 95% Bayesian credibility interval, and the solid line is the posterior median. Vertical dashed line is the mean TMRCA.

### Sources and frequencies of the transmissions between different hosts and farms

Host (humans and minks) and farm number labels were added to the sequences, and the number of transmissions between hosts (asymmetric) and between farms (symmetric) were inferred using discrete traits models on the time resolved trees (Figure 3, Figure S1 and S2). To avoid sample size effect on the results, sequences were further subsampled to reduce over-representative sequences from the same farm. For transmissions identified by Markov jumps, we also used BSSVS to identify only statistically significant pairs (with Bayes Factor >3, the higher the value, the stronger the support). We summarised and compared the network among three clusters A, C and D.

Overall, at least 43 zoonotic transmissions (with 95% HPD 34 to 50) from minks to humans likely occurred in multiple farms (Table S2). Specific, an average of 27 transmissions of viruses belonging to Cluster A occurred within 13 farms (NB1, NB3, NB8, NB13, NB21, NB52, NB55, NB56, NB57, NB58, NB59, NB63, NB68); 10 transmissions of virus belonging to Cluster C occurred within 7 farms (NB7, NB9, NB14, NB17, NB26, NB29, NB32) and 6 jumps of viruses belonging to Cluster D occurred within 3 farms (NB15, NB18 and NB19). However, some human infections may also be due to human-to-human infections, between mink farm employees or farm owner family members, which is not included in the model. Therefore, the true number of mink-to-human jumps may be lower.

There are also a few jumps between humans and minks from different farms. For example, within Cluster A, a sequence from humans linked to NB49 are likely transmitted from minks on NB47, although the low number of sequences (there is only one mink sequence obtained in NB49) precludes robust conclusions. We found that viruses may jump back and forth between humans and minks. The sequences sampled from humans in NB8 are likely transmitted to minks in NB12, as shown in the phylogeny of Cluster A (Figure 3). Epidemiology data indeed shows that the two farms have personnel links, which could be the explanation of this observation (supporting file 1).

We also identified different potential transmission patterns networks between farms in Cluster A, C and D (Figure 6 and Table S3). For Cluster A, NB47 seems to be the most important donor, with transmission to 7 farms (Figure 6a). In comparison, fewer significant between farm transmissions are identified in Cluster C and D (Figure 6b and c). Transmissions were also drawn as links between locations of mink farms on the map (Figure 7). Interestingly, we found transmissions with high BF supports (darker red edges in Figure 7a) that are not necessarily between adjacent farms, and sequences from adjacent farms with personnel links (e.g., NB58 and NB59). In addition, sequences from different barns on the same farms do not necessarily group together. For example, within Cluster C, sequences isolated from NB6 at the same date fell into two separate sub-clades.

**Figure 6.**
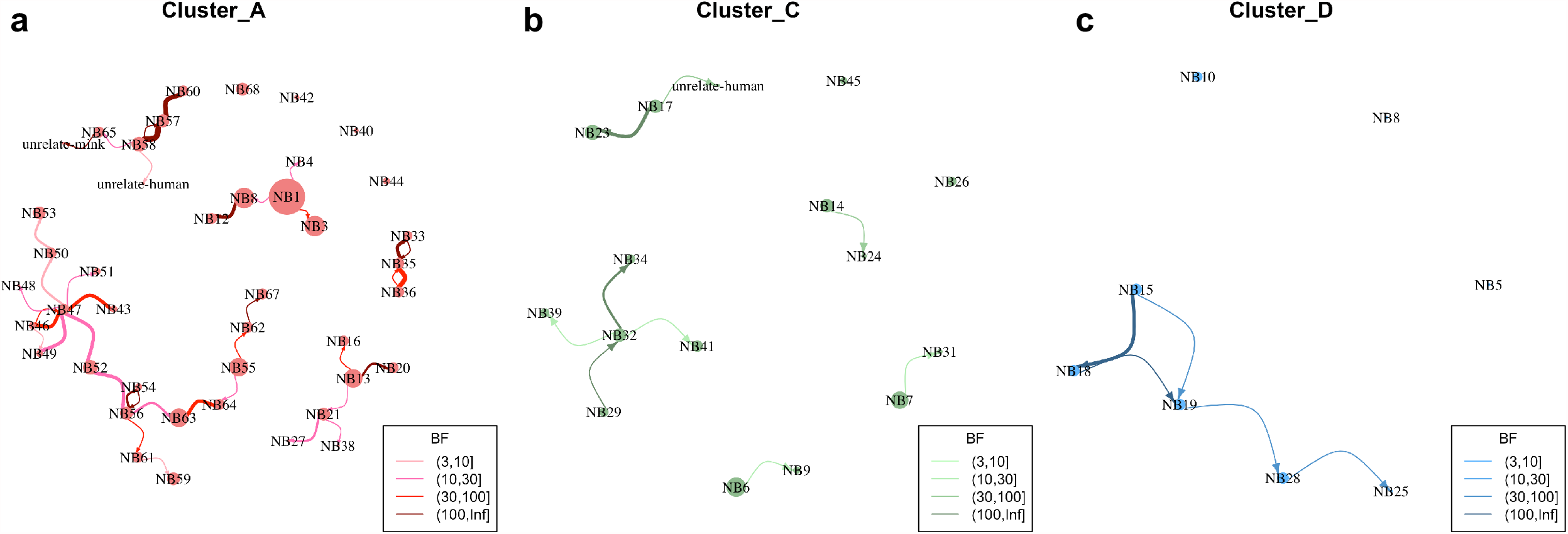
Transmission network between farms inferred from phylogenies of 3 mink clusters. **a** Cluster A **b** Cluster C and **c** Cluster D. Size of node indicates number of samples; edge weight indicates median number of transmissions between pairs of farms; arrow on edge indicates transmission direction; color of edge from light to dark indicates Bayes Factor (BF) support from low to high only transmissions with BF >3 are shown). The correlated farms are grouped together. Nodes with no link to the others indicated no significant transmissions with other farms although sequences belong to the cluster have been sampled. The number of transmissions and the correlated BF supports are shown in Table S3.

**Figure 7.**
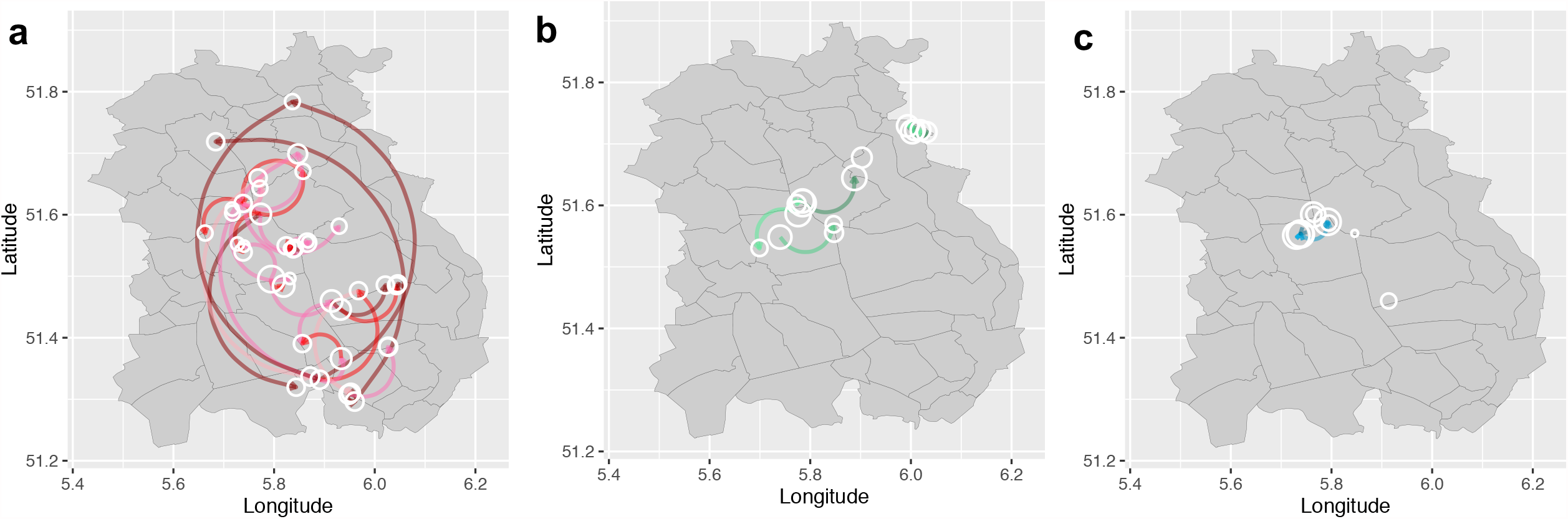
Transmission network between farms on map inferred from phylogenies of 3 mink clusters. **a** Cluster A **b** Cluster C and **c** Cluster D. The locations of farms on the map have been jittered for privacy reasons. Size of nodes indicates number of samples; arrow on edge indicates transmission direction; color of edge from light to dark indicates Bayes Factor support (BF) from low to high (only transmissions with BF >3 are shown), color keys are the same as Figure 6.

Assuming the presence of farm specific signatures allowed linking cases to farms, the two unrelated human sequences are most closely related to sequences from to farms NB17 and NB58, respectively; and the sequence from an escape mink is likely to have a relation with farm NB65 (Figure 6). However, the patients infected with mink strains did not report any direct or indirect contact with mink or mink farm employees.

### Inferred predictors of transmissions between farms

During our study, a detailed inventory of possible common characteristics, including farm owner, shared personnel, feed supplier and veterinary service provider was made. Epidemiological investigation indicated that many farms shared the same feed supplier or veterinarian, but no unambiguous service company contacts were found between farms within the different virus clusters which could explain the farm-to-farm spread. For 55% of the SARS-CoV-2 positive farms, owners, family members or personnel, including people with limited contact with minks, were shared between farms (supporting file 1).

Using a generalized linear model (GLM), implemented in BEAST, we tested the contribution of a range of predictor variables to the spread of viruses between farms which was estimated in the discrete trait phylogeographic model (Figure 6). Correlations between the predictor data collected from mink farms were tested and highly correlated predictors were omitted (Figure S4). The predictors being tested are 1) distance between farms; 2) personnel links between farms 3) feed supplier; 4) veterinary service provider; 5) mink population per farm; 6) number of sequences per farm included in the phylogenetic analysis; 7) human population density in municipality where farm was located; 8) days between sampling and culling per farm (supporting file 1).

For Cluster A, the distance between farms had a negative impact on the transmission between farms (Table 1), which indicated that farms that are further apart have generally lower rate of transmission between them; while farms with personnel links have a positive impact on the transmission between farms, which could be an explanation of the strong supported long-distance diffusion observed in Figure 7. For Cluster C and D, none of the predictors have significant impact on the overall transmission between farms (Table S4).

**Table 1.**
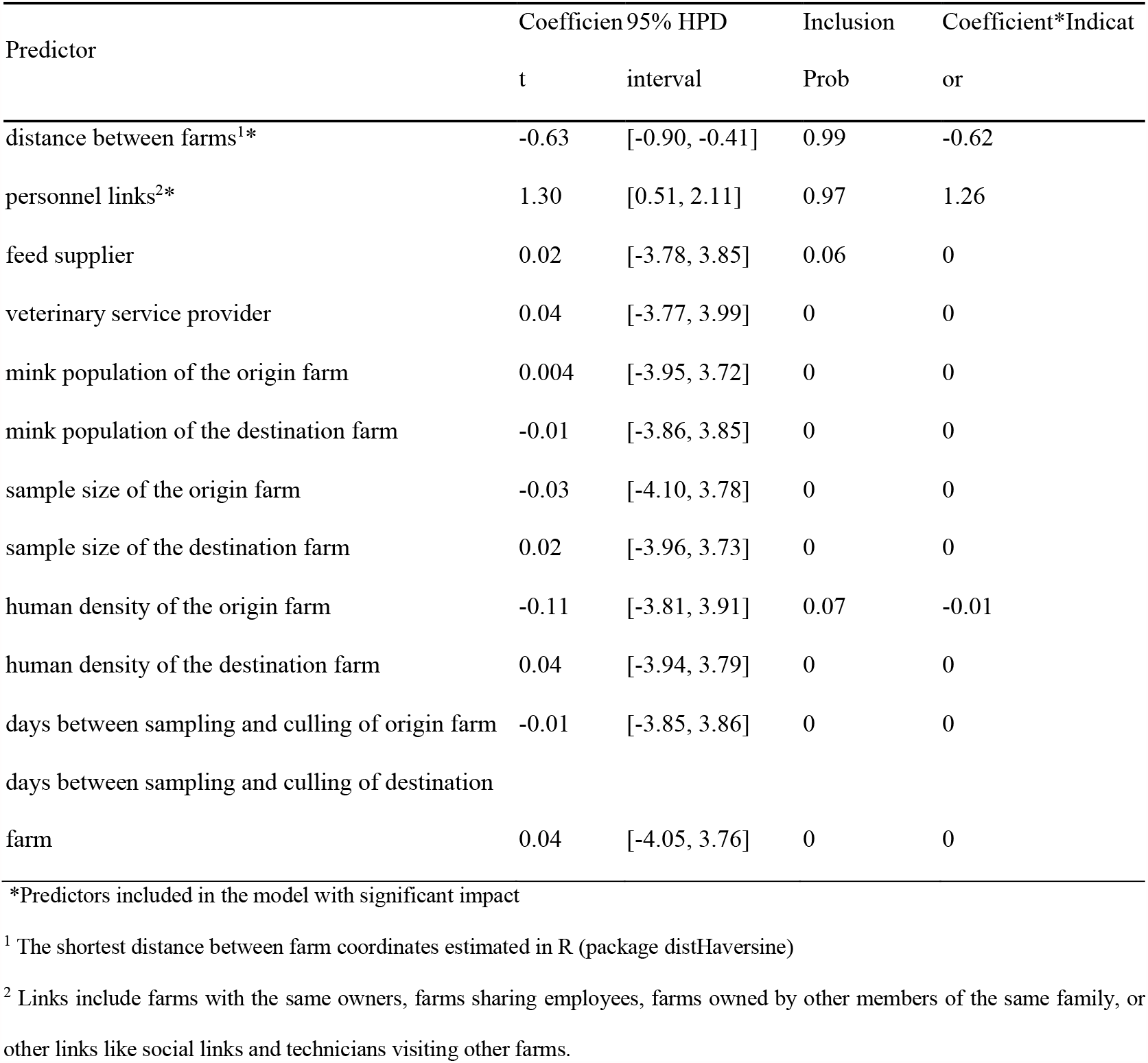
The contribution of predictors of SARS-CoV-2 (Cluster A) transmissions between mink farms

## Discussion

In this study, we explored the transmission dynamics of SARS-CoV-2, between mink farms, and between minks and humans, by combining SARS-CoV-2 monitoring in humans and animals, associated epidemiological information and the phylodynamic and transmission patterns of different SARS-CoV-2 sequence clusters in minks and in humans.

SARS-CoV-2 has infected 100 million people worldwide. Over 1,500,000 genomes have been generated and more than 800 lineages contributed to the active spread globally by 1^st^ April 2021 (when the analysis was performed) ^5^. Within the Netherlands, at least 140 lineages have been circulating in humans. We found five distinct clusters (A-E) derived from 4 different lineages (B.1.8, B.11, B.1.22, B.1.5) which had been dominantly circulating in the general human population in the Netherlands until 1^st^ April 2021. The most recent common ancestors of the five different mink clusters appeared in the Netherlands between mid-March to late-April 2020, which is in line with the timing of initial human detections in the country ^8^. The timing of introductions and expansions into mink populations are commensurate with exponential growth of SARS-CoV-2 in the human population in the Netherlands and with the mating season of the farmed minks, which is associated with an increase in use of external labour with more chances to have contact with humans ^3^. The last infected farm was detected in November 2020, after which no new infections were detected, probably due to lack of remaining farms with minks in the affected area and the start of the pelting season during which all minks, including the adults, were pelted due to the ban on mink farming from January 2021 onwards.

In comparison, the Cluster V variant, found in farmed minks in Denmark, is derived from a Danish specific lineage B.1.1.298 (https://cov-lineages.org/pango_lineages.html). This variant had 4 specific Spike mutations (69del, Y453F, I692V, M1229I), where Y453F was thought to be strongly associated with the mink infections and may be associated with decreased antibody binding and increased ACE2 affinity ^9-11^. Cluster V viruses were also found to infect humans and were associated with community transmission after mink-to-human transmission ^12^. Currently, viruses with the Y453F mutation have been identified in ∼1500 SARS-CoV-2 genomes and in 24 different lineages from Europe, Africa and the USA. Other potentially important mutations found in minks in Denmark in the spike protein (I692F, M1229I) can also be found in humans globally. Here we highlighted Y453F together with other 3 mutations F486L, 452M and N501T, which were first identified in multiple mink clusters that infected both farmed minks and related humans in the Netherlands. Their exact implications for viral fitness, transmissibility, and antigenicity need further investigation.

SARS-CoV-2 infections of minks are concerning as evolution of the virus in an animal reservoir could lead to establishment of additional zoonotic reservoirs with the potential for recurrent spill-over events of novel SARS-CoV-2 variants from minks to humans and other mammals ^13^. The spread of SARS-CoV-2 among farms was examined using phylodynamic methods. After the sudden increase in incidence of SARS-CoV-2 (mainly Cluster A) positive mink farms in August 2020 we observed that the virus had acquired several mutations compared to the virus last detected at the end of June, including the F486L mutation in spike protein. It is plausible that the increased phylodynamic growth rate (Re) after summer 2020 is associated with increased transmissibility in minks due to the emergence of clades with specific mutations in the spike protein ^14^. Interestingly, mutations at positions 452 and 501 have also been found emerging in some variants of interest ^10,11,15^. However, there is currently no evidence that these mutations are equivalent to the vital changes now seen in VOCs (Variants of Concern) in humans and cause a substantial shift in virus properties that enabling much better transmission in humans.

Our findings suggested that a personnel link is one key driver to explain the subsequent transmission among minks and transmission between different mink farms. Other factors than transmissions via humans are less likely to contribute in the cases where long distance transmissions occurred. Nevertheless, there was generally a positive association with farms in closer proximity, which is consistent with studies on SARS-CoV-2 infections in minks in other countries ^16^ and on other pathogens ^17-19^. There are also other potential drivers of between-farm transmissions of SARS-CoV-2 ^20,21^. For example, SARS-CoV-2 has been detected in feral cats and dogs around Dutch mink farms, showing evidence of mink-to-cat transmission of SARS-CoV-2 ^22,23^. In addition, free ranging mustelids have tested positive in other countries as well as two escaped mink in our study ^21^. Although for some other pathogens, farm-to-farm transmission via air has been proposed, SARS-CoV-2 RNA in ambient air outside of infected mink farms was not detected ^24^. The number of humans with mink strains around mink farms was nearly absent, making this scenario less likely as well.

We observed varied phylodynamic and transmission patterns among different mink clusters: the largest Cluster A emerged earlier and has comparably higher evolutionary rate and faster and wider spatial spread over a longer period of time than other clusters. However, for clusters for which we have fewer samples available, we observed higher uncertainty of the estimated phylodynamic parameters (e.g., Ne and Re with wide HPD intervals). In addition, the possibilities of missing samples in clusters would also lead to a putative bias in the trait analyses and GLM on identifying the significant transmission network and associated predictors and therefore we need to be cautious not to overinterpret the results. For example, the impact of humans on transmission between farms may still be underestimated as it is difficult to identify, locate and sample unregistered or moving workers in mink farms. Moreover, of 102 known human infections on mink farms, only 57 were successfully sequenced.

Finally, we identified multiple events of mink clusters jumping back and forth between human and minks within several mink farms. These infections were limited to people related to the farms with limited spread observed in the general population. However, the mink farming system and associated biosecurity policies may be different in other countries, possibly increasing risk of mink infections for humans. Moreover, with increasing human vaccination rate, as well as potentially animal vaccination, the relative importance and contribution to SARS-CoV-2 evolution of potential animal reservoirs may become more important. Although, the Cluster V variant was found in a substantial part of the population in Northern Jutland region of Denmark, the variant has not been detected anymore after November 2020, potentially due to culling of infected mink farms ^12,25^. This was also the case in the Netherlands all infected mink farms have been culled. The high number of infections in Dutch mink farms and associated human owners and workers, combined with the specific mutations found in the spike region and other regions of the SARS-CoV-2, shows that continuous surveillance and preventive measures in the fur farming industry ^16^, as well as other susceptible animal populations are advisable. Moreover, the emergence of novel variants may also have an effect on the virus’ host range, as has already been shown for the ability to infect mice of the Beta and Gamma variant, as opposed to the wildtype virus and the Alpha variant ^26^. Therefore, it is essential to keep monitoring the behaviour of the virus in combination with genetic information in both human and animals, especially animal species that have close contact with humans.

## Methods

### Samples and metadata

#### Mink

Mink farms suspected of SARS-CoV-2 infections were visited for sampling and epidemiological investigation by the competent authority (Netherlands Food and Consumer Product Safety Authority, NVWA). Farms were visited based on reporting of increased mortality or respiratory signs by owners or when tested positive during surveillance systems (Figure 1a). These included an Early Warning system (EWS) of weekly testing of carcasses of recently dead minks by RT-PCR on throat samples and mandatory serological screening by GD Animal Health (GD, Deventer, the Netherlands) ^16^.

Official sampling included non-random sampling of 20 minks, by means of throat and rectal swabs, targeting minks with clinical signs. Throat swabs of two minks, caught at the end of September / beginning of October 8 and 9 days after culling of two farms (NB58 and NB59) at 450 and 650 m distance respectively, which most likely escaped during culling, were also submitted for testing. Associated metadata was derived from the database developed by a consortium of One Health outbreak experts. Data collected for each farm included farm location, number of animals, ownership, shared personnel and other contacts (anonymised), data of confirmed SARS-CoV-2 detection and time interval between sampling and culling. The epidemiology data are in supporting file 1.

#### Human cases related to mink farms

On the first SARS-CoV-2 infected mink farms, NB1-NB16 (NB is the Dutch abbreviation for mink farm, which were numbered consecutively based on diagnosis) active case finding as well as serum collection of people with possible exposure to infected minks was performed, as described previously ^4^. On farms NB17-NB68, all owners and employees of infected mink farms were requested to visit a regional SARS-CoV-2 testing facility in case of any symptoms indicative of COVID-19, in line with the national SARS-CoV-2 testing and surveillance policy. There were no serum samples taken for antibody detection.

#### Medical ethical permission

Outbreak investigations of notifiable diseases such as COVID-19 are the legal tasks of the Public Health Service as described under the Public Health Act, and do not require separate medical ethical clearance.

#### 4-digit post code screening

Two screenings of SARS-CoV-2 positive humans living in the same region as the infected mink farms took place from 3^rd^ April 2020 to 16^th^ November 2020. The first screening included a set of sequences obtained from anonymized samples from patients that had been diagnosed with COVID-19 in the area of the same four-digit postal codes as farms NB1-NB4 in March and April 2020, as described previously ^4^. For the second screening, municipal health centres selected anonymised laboratory IDs for 10 SARS-CoV-2 positive humans in the period 15^th^ October 2020 to 16^th^ November 2020 from the same postal code regions of the 68 SARS-CoV-2 positive mink farms from their notification system. Based on the laboratory ID, stored samples were retrieved from the diagnostic centres for sequencing. In some regions the number of samples was lower than 10, due to low numbers of positives in the selected period or because not all samples had been retained by the laboratories. Samples from the selected postal codes that were collected in the period 27^th^ November 2020 to 4^th^ January 2021 were also included in the analysis.

#### SARS-CoV-2 diagnostics and sequencing

Human and animal cases were diagnosed by SARS-CoV-2 RT-PCR testing of oropharyngeal and rectal swabs (minks) or upper respiratory tract samples (humans) in one of the laboratories participating in the national COVID-19 response ^27^. RT-PCR positive samples were processed for whole genome sequencing as described previously ^4^. For each mink farm, a maximum of five of the RT-PCR positive samples with Ct<32 were selected, based on lowest Ct-values.

For the mandatory serological screening in mink, blood on filter paper was eluated and approximately 2 µL of serum was tested for SARS-CoV-2 antibodies using an in-house indirect ELISA based on the RBD antigen. The same ELISA using the S1 antigen was used for confirmation (unpublished).

The first and last 30 nucleotides were trimmed, and subsequently mapped against the NC_045512.2 SARS-CoV-2 reference genome using minimap2 ^28^. After mapping the alignment files were used to generate a consensus sequence using pysam module ^29^ in a custom python script. Homopolymeric regions were manually checked and resolved by consulting reference genomes and positions with less than 30x coverage were replaced with “N” ^30^. The complete sequences information and metadata used in the phylogenetic analyses are in supporting file 2.

### Phylodynamic reconstructions

Complete SARS-CoV-2 genomes with >95% coverage isolated from minks and related humans were included in the phylodynamic reconstructions. We also included human sequences from across the Netherlands as background data. The data were obtained from GISAID (https://www.gisaid.org/) and the collected date was up to 4^th^ January 2021. We then subsampled these background human sequences to keep at least 1 sequence per global lineage as defined using the Pango-lineage classification (version 1^st^ April 2021) ^5^ per region per week.

Genomes were aligned with MAFFT ^30^ and edited by partitioning into coding regions and non-coding intergenic regions with a final alignment length of 29,508 nucleotides. Phylogenetic trees were first generated using IQtree^31^ employing maximum likelihood (ML) under 1000 bootstraps. The nucleotide substitution model used for all phylogenetic analyses was HKY with a Gamma rate heterogeneity among sites with four rate categories. To determine if our sequence data exhibited temporal qualities, we used TempEst v1.5 ^32^ to measure the root-to-tip divergence for ML trees.

Phylodynamic analyses of SARS-CoV-2 in mink farms in Netherlands were conducted using time-scaled Bayesian phylogenetic methods in BEAST version 1.10.4 ^33^. The best fit models were HKY+G+4 for the site substitution model and skygrid ^34^ for the tree model, determined by using stepping-stone sampling^35^. We first generated phylogeny using all full-length genomes of SARS-CoV-2 from mink farms with background human samples using an uncorrelated relaxed molecular clock model which assumes each branch has its own independent substitution rate ^36^, We then generated independent phylogenies of Cluster A, C and D using a strict molecular clock model with prior specified (a mean of 1×10-3 with 95% HPD between 6×10^−4^ and 2×10^−^ 3). To analyse fluctuations in SARS-COV-2 epidemic spread in mink farms in the Netherlands per individual cluster, we estimated the changes of viral effective population size (Ne) over time using the skygrid model ^34^ in BEAST version 1.10.4, and the effective reproductive number (Re) during the course of the outbreak in mink farms, using the Birth-death skyline (BDSKY) model ^37^ in BEAST2 version 2.6.3 ^38^. The effective reproductive number (Re) is estimated from the time-scaled phylogeny as a version of the phylodynamic lineage growth rate, and is representative of a between farm Re. We also used the multitype-tree birth-death model (BDMM) to explore whether the appearances of certain mutations in the spike protein have impact on the Re variations ^39^. We specified the following priors according to the knowledge of SARS-CoV-2 infections in humans and our epidemiology surveillance data on mink farm infections: 1) Re: a mean of R0 2.5 with 95% HPD (0.6, 6) ^20,40^, and were estimated over 5 equidistant time intervals depending on the size of the overall tree; 2) the “becomeUninfectiousRate”, which refers to the number of days from infection to culling for a mink/farm: a mean of 26 (equivalent to 14 days) with 95% HPD between 5 to 20 days; 3) the sampling portion, which refers to the number of sequences per farm divided by the total infected mink population of a farm: a mean of 2×10^−4^ with 95% HPD (1×10^−5^, 1×10^−3^) and 4) the origin time of the epidemic: the estimated time to the most recent common ancestors (TMRCAs) of the three mink clusters under strict clock model with priors described above. In addition, we compared the spatial diffusion rates (geography.clock.rate) among the 3 clusters using the coordinates of each infected mink farm via a continuous model in BEAST2 version For each analysis the MCMC algorithm was run for 10^8^ steps and sampled every 10^4^ steps.

We further estimated the transmissions between farms and between minks and humans using the phylogenies of Cluster A, C and D separately. We used an asymmetric model and incorporated BSSVS to identify a sparse set of transmission rates that identify the statistically supported connectivity ^41^. We also estimated the expected number of transmissions (jumps) between farms and hosts using Markov rewards ^42^. Finally, we inferred the possible predictors that may drive to the spread of virus between farms (estimated between-farm transmission rates) using a generalized linear model (GLM), an extension of the discrete diffusion model ^43^.

## Data availability

The sequence data and epidemiology data used in these analyses are available in supplementary file 1 and 2.

## Acknowledgements

We acknowledge the authors, originating and submitting laboratories of the sequences from GISAID’s EpiCov Database on which the phylogenetic analysis was based (see Supplement). All submitters of data may be contacted directly via the GISAID website www.gisaid.org.

This work is supported by European Union’s Horizon 2020 research and innovation programme under Grant No. 874735 (VEO). The mink farm outbreak investigation was funded by the Dutch Ministries of Health, Welfare and Sport, and of Agriculture, Nature and Food Quality.

We declare that we have no competing financial, professional, or personal interests that might have influenced the performance or presentation of the work described in this article.

## Author Contributions

LL, RSS and FCV wrote the manuscript. FCV, PAM, NBV, AR, MCAWB, WB, PT, MK, LAMS, RWHH, WHMP, ANS, MAHS RJM, JR, JAS and MA set up sample and data collection. BBOM, RSS, DFN generated sequence data. LL, RSS, FCV and EAJF were involved in data analysis and interpretation. LL, RSS, SL, BBOM, MPGK designed the study. All authors provided critical feedback and contributed to manuscript editing.

## Competing Interests

The authors declare no competing interests

## Notes

### Competing Interest Statement

The authors have declared no competing interest.

