## Supplementary figures and tables for "Adaptation, spread and transmission of SARS-CoV-2 in farmed minks and related humans in the Netherlands"

### Supplementary tables and figures

Table S1 Estimated TMRCA, evolution rate and spatial diffusion rate of Cluster A, C and D

| Cluster | TMRCA | | Last sampling date | Evolution.rate x10-3 | | Diffusion.rate x10-4 | |
|---|---|---|---|---|---|---|---|
| | mean | 95% HPD | | mean | 95% HPD | mean | 95% HPD |
| A | 21/03/2020 | [10/03/2020, 10/04/2020] | 04/11/2020 | 1.41 | [1.20, 1.75] | 2.91 | [2.45, 3.38] |
| C | 28/03/2020 | [12/03/2020, 22/04/2020] | 03/09/2020 | 0.88 | [0.57, 1.04] | 1.06 | [0.81, 1.34] |
| D | 19/04/2020 | [27/03/2020, 11/05/2020] | 08/08/2020 | 1.16 | [0.77, 1.60] | 1.34 | [0.61, 2.2] |

Table S2 Number of significant (BF>3) transmissions between human and minks of Cluster A, C and D

| Cluster | host | BF2 | median_jump | 95% HPD interval |
|---|---|---|---|---|
| A | human->mink | >100 | 4 | (0, 6) |
| | mink->human | >100 | 27 | (22, 32) |
| C | human->mink | >100 | 1 | (0, 1) |
| | mink->human | >100 | 10 | (7, 12) |
| D | human->mink | >100 | 0 | (0, 1) |
| | mink->human | >100 | 6 | (5, 6) |

Table S3 Number of significant (BF>3) transmissions between farms

| Cluster | farm | BF | median_jump | 95% HPD interval |
|---|---|---|---|---|
| A | NB1->NB3 | (30,100] | 1 | (0, 2) |
| | NB1->NB4 | (10,30] | 1 | (0, 2) |
| | NB1->NB8 | (10,30] | 1 | (0, 2) |
| | NB13->NB16 | (30,100] | 1 | (0, 2) |
| | NB13->NB20 | (100,Inf] | 2 | (1, 4) |
| | NB13->NB21 | (10,30] | 1 | (0, 2) |
| | NB21->NB27 | (10,30] | 1 | (0, 2) |
| | NB21->NB38 | (10,30] | 1 | (0, 2) |
| | NB33->NB35 | (100,Inf] | 1 | (0, 5) |
| | NB35->NB33 | (100,Inf] | 3 | (0, 6) |
| | NB35->NB36 | (30,100] | 4 | (0, 7) |
| | NB36->NB35 | (30,100] | 1 | (0, 4) |
| | NB46->NB47 | (30,100] | 1 | (0, 7) |
| | NB47->NB43 | (30,100] | 3 | (0, 5) |
| | NB47->NB46 | (30,100] | 3 | (0, 6) |
| | NB47->NB48 | (10,30] | 1 | (0, 2) |
| | NB47->NB49 | (10,30] | 3 | (0, 6) |
| | NB47->NB50 | (3,10] | 2 | (0, 3) |
| | NB47->NB51 | (10,30] | 1 | (0, 2) |
| | NB47->NB52 | (10,30] | 3 | (0, 7) |
| | NB50->NB53 | (3,10] | 2 | (1, 3) |
| | NB52->NB56 | (10,30] | 3 | (0, 6) |
| | NB54->NB56 | (100,Inf] | 1 | (0, 5) |
| | NB55->NB62 | (30,100] | 1 | (0, 2) |
| | NB55->NB64 | (10,30] | 1 | (0, 2) |

| | | | | | |
|---|---|---|---|---|---|
| | NB56->NB54 | (100,Inf] | 2 | | (0, 4) |
| | NB56->NB61 | (30,100] | 1 | | (0, 4) |
| | NB56->NB63 | (10,30] | 2 | | (0, 6) |
| | NB57->NB58 | (100,Inf] | 5 | | (0, 7) |
| | NB57->NB60 | (100,Inf] | 4 | | (0, 6) |
| | NB58->NB57 | (100,Inf] | 1 | | (0, 5) |
| | NB58->NB65 | (10,30] | 1 | | (0, 2) |
| | NB58->unrelate-human | (3,10] | 1 | | (0, 2) |
| | NB61->NB59 | (3,10] | 1 | | (0, 4) |
| | NB62->NB67 | (100,Inf] | 1 | | (1, 2) |
| | NB63->NB64 | (30,100] | 3 | | (0, 5) |
| | NB65->unrelate-mink | (100,Inf] | 1 | | (1, 1) |
| | NB8->NB12 | (100,Inf] | | 3 | (1, 5) |
| C | NB14->NB24 | (10,30] | 1 | | (0, 1) |
| | NB17->NB23 | (100,Inf] | 3 | | (1, 5) |
| | NB17->unrelate-human | (30,100] | 1 | | (0, 1) |
| | NB7->NB31 | (10,30] | 1 | | (0, 1) |
| | NB29->NB32 | (100,Inf] | 1 | | (0, 5) |
| | NB32->NB34 | (100,Inf] | 2 | | (0, 5) |
| | NB32->NB39 | (10,30] | 1 | | (0, 4) |
| | NB32->NB41 | (10,30] | 1 | | (0, 2) |
| | NB6->NB9 | (10,30] | 1 | | (0, 3) |
| D | NB15->NB18 | >100 | 3 | | (0, 3) |
| | NB15->NB19 | (3,10] | 1 | | (0, 2) |
| | NB18->NB19 | >100 | 1 | | (0, 2) |
| | NB19->NB25 | (30,100] | 1 | | (0, 2) |

| | NB25->NB28 | (30,100] | 1 | (0, 2) |
|---|---|---|---|---|

Table S4 The contribution of predictors of Cluster A, C and D transmissions between mink farms

| Cluster | Predictor | Coefficient | 95% HPD interval | Inclusion Prob | Coefficient*Indicator |
|---|---|---|---|---|---|
| A | distance between farms* | -0.63 | [-0.90, -0.41] | 0.99 | -0.62 |
| | personnel links* | 1.30 | [0.51, 2.11] | 0.97 | 1.26 |
| | feed supplier | 0.02 | [-3.78, 3.85] | 0.06 | 0 |
| | veterinary service provider | 0.04 | [-3.77, 3.99] | 0 | 0 |
| | mink population of the origin | 0.004 | [-3.95, 3.72] | 0 | 0 |
| | mink population of the destination | -0.01 | [-3.86, 3.85] | 0 | 0 |
| | sample size of the origin | -0.03 | [-4.10, 3.78] | 0 | 0 |
| | sample size of the destination | 0.02 | [-3.96, 3.73] | 0 | 0 |
| | human density of origin | -0.11 | [-3.81, 3.91] | 0.07 | -0.01 |
| | human density of destination | 0.04 | [-3.94, 3.79] | 0 | 0 |
| | days between sampling and culling origin | -0.01 | [-3.85, 3.86] | 0 | 0 |
| | days between sampling and culling destination | 0.04 | [-4.05, 3.76] | 0 | 0 |
| C | distance between farms | -0.24 | [-3.45, 3.67] | 0.38 | -0.09 |
| | personnel links | 0.01 | [-3.79, 3.86] | 0.04 | 0 |
| | feed supplier | -0.01 | [-4.01, 3.85] | 0.01 | 0 |
| | veterinary service provider | 0.02 | [-3.855, 3.9941] | 0.02 | 0 |
| | mink population of the origin | 0.08 | [-4.08, 3.87] | 0.03 | 0.002 |
| | mink population of the destination | -0.01 | [-3.84, 3.94] | 0.06 | -0.001 |
| | sample size of the origin | -0.02 | [-3.88, 3.76] | 0.01 | 0 |
| | sample size of the destination | -0.01 | [-3.96, 3.88] | 0.01 | 0 |

| | | | | | |
|---|---|---|---|---|---|
| | human density of origin | 0.03 | [-3.93, 3.87] | 0.01 | 0 |
| | human density of destination | -0.002 | [-3.89, 3.97] | 0.01 | 0 |
| | days between sampling and culling origin | 0.10 | [-3.76, 3.88] | 0 | 0 |
| | days between sampling and culling destination | -0.01 | [-3.77, 4.08] | 0 | 0 |
| D | distance between farms | 0.01 | [-3.77, 3.92] | 0.01 | 0 |
| | personnel links | 0.02 | [-4.04, 3.76] | 0.01 | 0 |
| | feed supplier | 0 | [-4.01, 3.81] | 0 | 0 |
| | veterinary service provider | -0.02 | [-3.72, 3.99] | 0 | 0 |
| | mink population of the origin | 0.02 | [-3.54, 4.17] | 0 | 0 |
| | mink population of the destination | 0.01 | [-4.07, 3.75] | 0 | 0 |
| | sample size of the origin | 0.20 | [-3.86, 4.02] | 0.07 | 0.014 |
| | sample size of the destination | -0.01 | [-3.89, 3.74] | 0.00 | 0 |
| | human density of origin | 0.12 | [-3.78, 4.05] | 0.06 | 0.01 |
| | human density of destination | -0.01 | [-3.74, 4.03] | 0.01 | 0 |
| | days between sampling and culling origin | 0.03 | [-3.90, 3.82] | 0.01 | 0 |
| | days between sampling and culling destination | -0.01 | [-3.89, 3.80] | 0 | 0 |

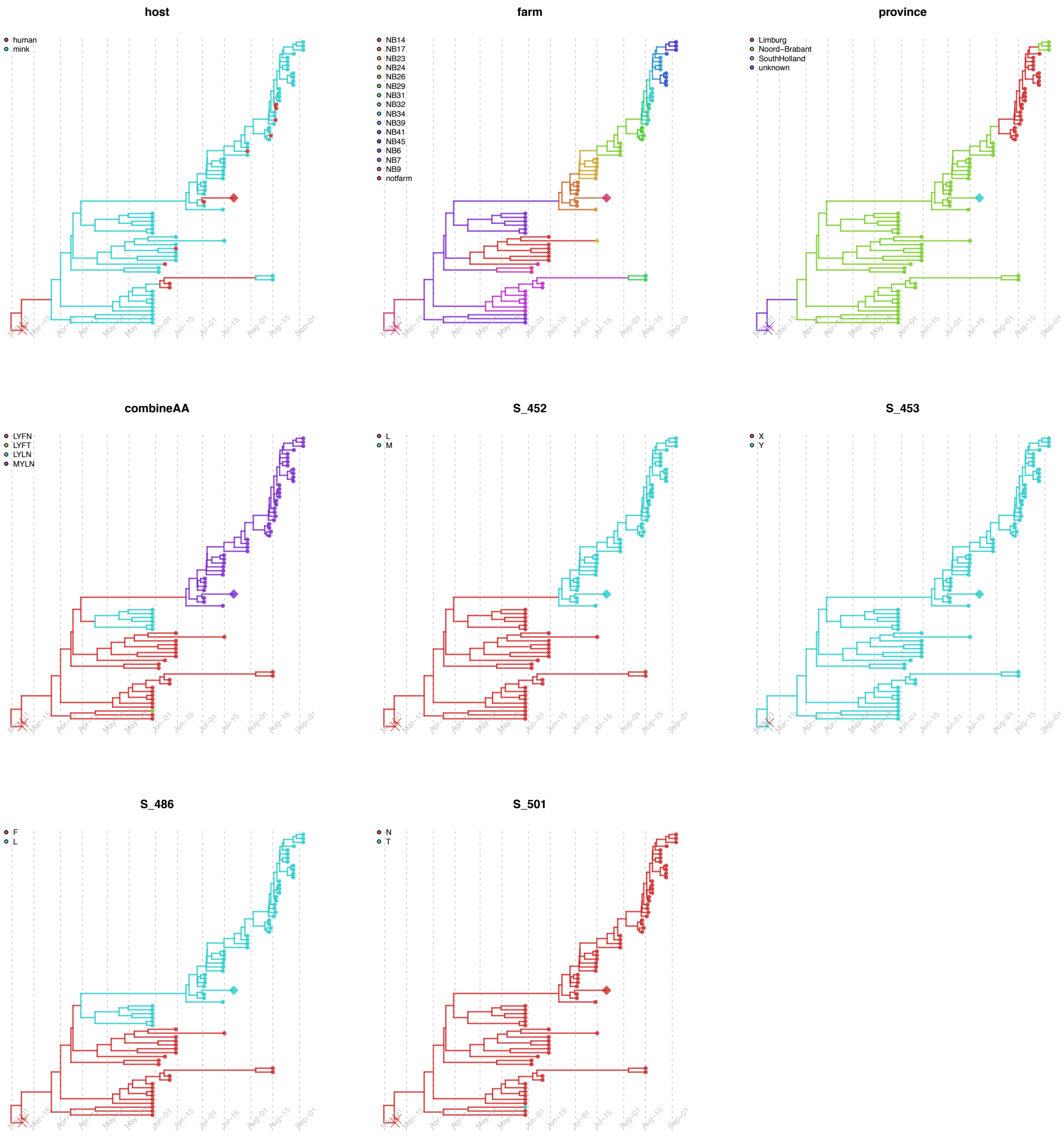


Figure S1. Discrete trait mapping on time-scaled phylogeny of Cluster C. The 8 types of traits are: host, farm number, province, the 4 selected mutations on spike protein and the combinations of the 4 mutations. The branches and nodes are colored by inferred ancestral traits. One unrelated human sequence fell within mink clusters but not isolated from farms is highlighted in diamond. The outgroup human samples in the origin are cross labelled.

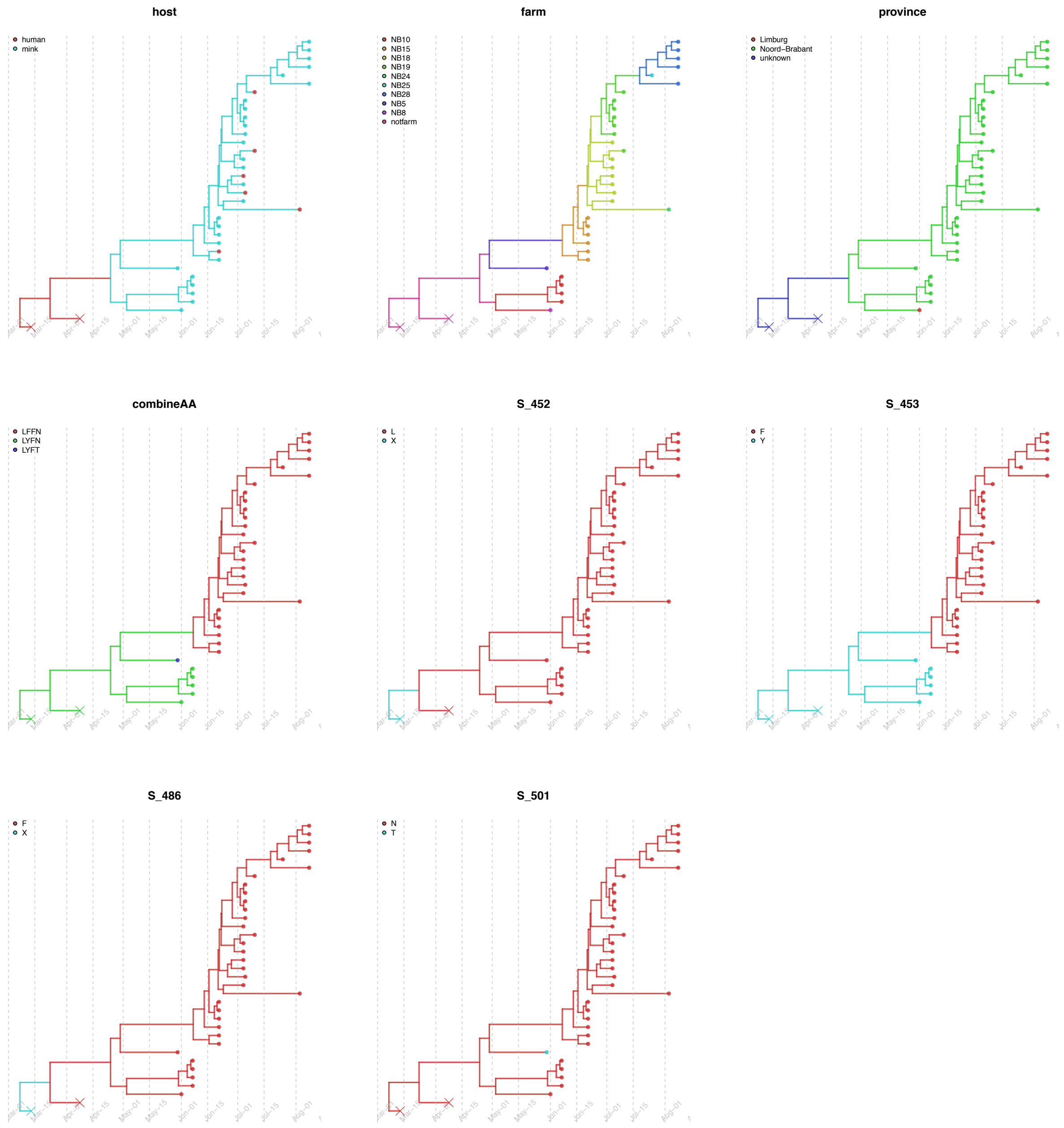


Figure S2. Discrete trait mapping on time-scaled phylogeny of Cluster D. The 8 types of traits are: host, farm number, province, the 4 selected mutations on spike protein and the combinations of the 4 mutations. The branches and nodes are colored by inferred ancestral traits. The outgroup human samples in the origin are cross labelled.

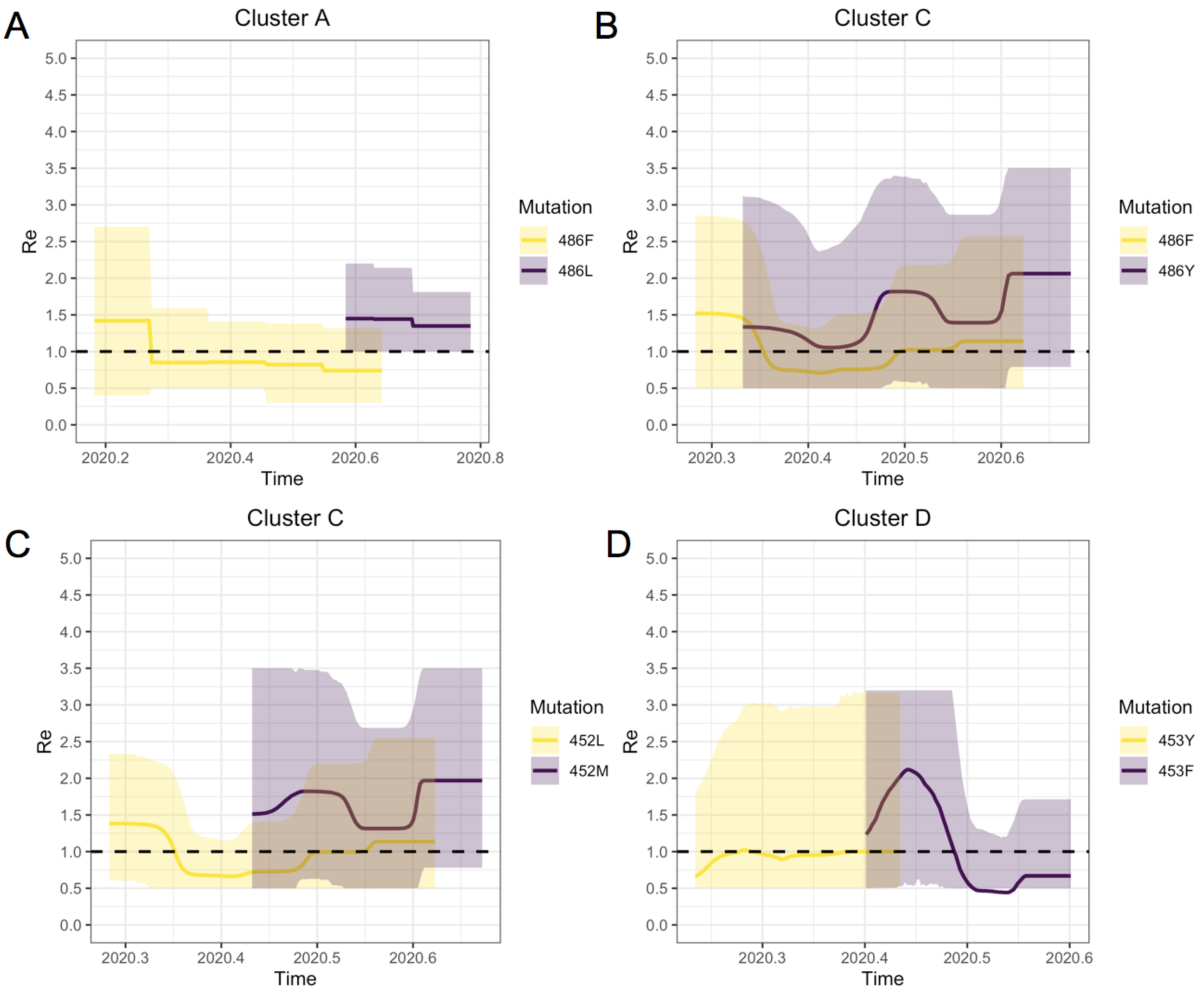


Figure S3 Multitype birth-death (BDMM) analysis of impact of appearance of mutations on Spike on reproductive number over time. For Cluster A, C and D, the changes in the Re's for clades with mutations on Spike protein site 486, 452 and 453 are shown in purple, clades without mutations are shown in yellow.

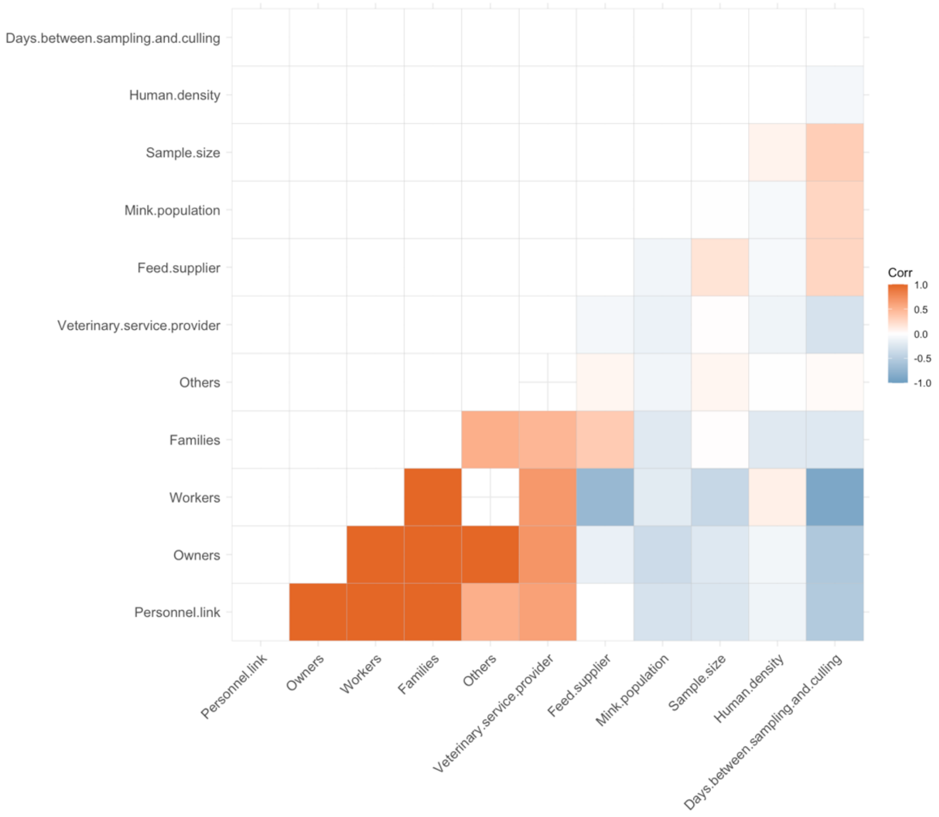


Figure S4 Correlation matrix between the predictor data collected from mink farms. The color corresponding to the correlation coefficient. Dark blue indicates strong positive correlation whereas dark red indicates strong negative correlation. Highly correlated predictors (sharing owners, sharing employees, owned by other members of the same family, or other links) were omitted.

#### Supporting files

Supporting file 1. The epidemiology metadata of mink farm NB1-68.

Supporting file 2. SARS-Cov-2 sequences isolated from minks and related-humans used in the phylogenetic analysis
